## Supplemental File for "Polymeric nature of tandemly repeated genes enhances assembly of constitutive heterochromatin in fission yeast"

### Supplementary Appendix: Connectivity of tandemly repeated genes enhances assembly of constitutive heterochromatin in fission yeast

Tetsuya Yamamoto,[†] Takahiro Asanuma,[‡] and Yota Murakami[‡]

†*Institute for Chemical Design and Discovery, Hokkaido University*

‡*Department of Chemistry, Faculty of Science, Hokkaido University*

### S1 Surface adhesion of polymers

We here discuss the essence of the adhesion of a polymer to a surface. To this end, we treat a section in a long polymer chain at the vicinity of a surface, see fig. S1**a**. The polymer section is composed of $N$ units that are adhesive to the surface. The section is connected to the surface via a linker chain composed of $N_{\rm L}$ units so that the adhesive section does not diffuse away from the surface ($N \ll N_{\rm L}$). The length of both units is $b$. We represent the positions in the system by the distance $z$ from the surface. Because the system is symmetric with respect to the $x$- and $y$- directions, we only analyze the motion of beads in the $z$-direction. Each adhesive unit can bind to the surface with the rate $k_{\rm on}$ when it is located in $0 < z < b$. A bound unit is unbound from the surface with the rate $k_{\rm off}$.

#### S1.1 Polymer with one adhesive unit

One can solve derive the analytic form of the binding probability $p$ for the case of a polymer that has only one adhesive unit, $N = 1$. The distribution of the adhesive unit is

$$\psi(z) = \frac{3z}{b^2 N_{\rm L}} {\rm e}^{-3z^2/(2b^2 N_{\rm L})}. \tag{S1}$$

The probability with which the adhesive unit is located in the region $0 < z < b$ is

$$\Psi = \int_0^b dz\, \psi(z) = 1 - {\rm e}^{-3/(2N_{\rm L})} \simeq \frac{3}{2N_{\rm L}}. \tag{S2}$$

The kinetic equation of the binding and unbinding of the adhesive unit has the form

$$\frac{dp}{dt} = k_{\rm on} \frac{3}{2N_{\rm L}} (1 - p) - k_{\rm off} p. \tag{S3}$$

In the steady state, $dp/dt = 0$, the binding probability is derived as

$$p = \frac{1}{1 + \frac{2}{3}\frac{k_{\rm off}}{k_{\rm on}} N_{\rm L}}. \tag{S4}$$

#### S1.2 Rouse dynamics simulation

We performed the Rouse dynamics simulation with which a polymer chain is modeled as beads connected by springs. The polymer chain is end-grafted to a surface and the beads are labeled as $n = 0, 2, \cdots, N + N_{\rm L}$ from the grafted bead. The equation of motion of the $n$-th bead (Langevin equation) has the form

$$\frac{d}{dt} z_n(t) = \frac{p_n}{m} \tag{S5}$$

$$\frac{d}{dt} p_n(t) = -\frac{\partial}{\partial z_n} U - \xi p_n(t) + f_n(t) \tag{S6}$$

with $n = 0, 2, \cdots, N + N_{\rm L}$. $p_n(t)$ is the momentum of the $n$-th bead and $z_n(t)$ is the position (the distance from the surface) of the $n$-th bead. These are functions of time $t$. $\xi$ is the friction constant of a bead and $m$ is the mass of a bead. Eq. (S5) is the definition of the momentum and eq. (S6) is the Newton's equation of motion with the potential force (the first term), the friction force (the second term), and the random force $f_n(t)$ due to the thermal fluctuation (the third term). The random force follows the Gaussian statistics with the mean and correlation

$$\langle f_n(t) \rangle = 0 \tag{S7}$$

$$\langle f_n(t) f_m(t') \rangle = 2\zeta k_{\rm B} T \delta_{mn} \delta(t - t'), \tag{S8}$$

where $m$ and $n$ $(= 1, 2, \cdots, N + N_{\rm L})$ are indices of monomers and $\zeta$ $(= m\xi)$ is the rescaled friction constant. In general, $U$ includes the short-range interaction due to the connectivity of the beads, the long-range interaction due to the excluded volume between the beads, and the interaction between the beads and the surface. In our simulation, we neglect the long-range interaction (Rouse model). The potential function $U$ has the form

$$U = \sum_{n=0}^{N+N_\mathrm{L}} U_\mathrm{s}(z_n) + \frac{1}{2}k \sum_{n=1}^{N+N_\mathrm{L}} (z_n - z_{n-1})^2. \tag{S9}$$

with

$$U_\mathrm{s}(z) = \begin{cases} U_0 + U_0 \left( \frac{z_\mathrm{m}^{12}}{z^{12}} - \frac{2z_\mathrm{m}^6}{z^6} \right) & 0 < z < z_\mathrm{m} \\ 0 & z_\mathrm{m} < z \end{cases} \tag{S10}$$

The form of eq. (S10) is the simple Lennard-Jones potential, from which the attractive part is omitted.

We integrate eqs. (S5) and (S6) numerically by using the velocity Verlet algorithm.[1] The differential equations are approximated by using the finite difference method

$$z_n(t+\Delta t) = z_n(t) + c_1 v(t)\Delta t + c_2 a(t)\Delta t^2 + \delta r_n \tag{S11}$$

$$v_n(t+\Delta t) = c_0 v_n(t) + (c_1 - c_2)a(t)\Delta t + c_2 a(t+\Delta t)\Delta t + \delta v_n. \tag{S12}$$

with

$$c_0 = \mathrm{e}^{-\xi\Delta t} \tag{S13}$$

$$c_1 = \frac{1}{\xi\Delta t}\left(1 - \mathrm{e}^{-\xi\Delta t}\right) \tag{S14}$$

$$c_2 = \frac{1}{(\xi\Delta t)^2}\left(\xi\Delta t - 1 + \mathrm{e}^{-\xi\Delta t}\right). \tag{S15}$$

$v_n(t)$ $(= p_n(t)/m)$ is the velocity and $a_n(t)$ $(= -\partial U/\partial z_n)$ is the acceleration. In the numerical calculation, the positions $z_n(t)$ are rescaled by the length scale

$$l_\mathrm{s}^2 = \frac{k_\mathrm{B}T}{m}\xi^{-2}. \tag{S16}$$

and the velocity is rescaled by the scale

$$v_{\mathrm{s}} = \xi l_{\mathrm{s}}. \tag{S17}$$

The relaxation time of a monomer is

$$\tau = \frac{m\xi}{k}. \tag{S18}$$

The random values, $\delta r_n$ and $\delta v_n$, follow the Gaussian distribution

$$P(\delta r_n, \delta v_n) = \frac{1}{2\pi\sigma_z\sigma_v\sqrt{1-c_{zv}^2}} \mathrm{e}^{-\frac{1}{2(1-c_{zv}^2)}\left(\frac{\delta z_n^2}{\sigma_z^2}+\frac{\delta v_n^2}{\sigma_v^2}-2c_{zv}\frac{\delta z_n}{\sigma_z}\frac{\delta v_n}{\sigma_v}\right)} \tag{S19}$$

with

$$\sigma_z^2 = \langle \delta z_n^2 \rangle = l_{\mathrm{s}}^2\left(2\xi\Delta t - 3 + 4\mathrm{e}^{-\xi\Delta t} - \mathrm{e}^{-2\xi\Delta t}\right) \tag{S20}$$

$$\sigma_v^2 = \langle \delta v_n^2 \rangle = v_{\mathrm{s}}^2\left(1 - \mathrm{e}^{-2\xi\Delta t}\right) \tag{S21}$$

$$\sigma_z\sigma_v c_{zv} = \langle \delta z_n \delta v_n \rangle = l_{\mathrm{s}} v_{\mathrm{s}}\left(1 - \mathrm{e}^{-\xi\Delta t}\right)^2 \tag{S22}$$

The subchain $N_{\mathrm{L}} < n < N + N_{\mathrm{L}}$ is composed of adhesive units, while the subchain $0 < n < N_{\mathrm{L}}$ is a linker. An adhesive unit can bind to the surface with a rate $k_{\mathrm{on}}$ when it is located in $0 < z < b$. A bound adhesive unit can unbind from the surface with a rate $k_{\mathrm{off}}$. We used the mersenne twister to generate random numbers.

The initial positions of the beads are randomly generated. If the randomly generated position $z$ of a bead is smaller than $z_{\mathrm{m}}$, it is inverted to $z_n(0) = 2z_{\mathrm{m}} - z$. The system is equilibrated for time $t_{\mathrm{eq}}$ and then performed the simulation for time $t$. The ratio $p_{\mathrm{on}}$ of bound units and the probability $q_{\mathrm{on}}$ that more than one unit is bound to the surface are extracted. This simulation was performed for $M$ times with different initial conditions. The values of parameters used for the simulation are summarized in Table S1. Our simulations predict the binding probability for the case of a polymer with one adhesive unit in agreement with eq. (S4), see the cyan broken line and cyan dots in fig. S2. In this case, the adhesive unit is bound to the surface at any time if and only if the binding constant is zero.

#### S1.3 Self-consistent scheme for a very long chain

To treat the surface adhesion of polymers analytically, we consider the case in which the adhesive unit is very long so that the end effect of the section is negligible. In this case, the probability $p_\infty$ that a unit is bound to the surface does not depend on the choice of units and more than one unit is bound to the surface at any time. The problem that we are going to solve is basically the same as ref. [2] We focus on the $l$-th unit in the adhesive section. We consider the case in which the units between the $1-m+1$-th and $l+n-1$-th units are not bound to the surface and the $l-m$-th and $l+n$-th units are bound to the surface, see fig. S1**b**. The distribution of the $l$-th unit has the form

$$\psi_{m,n}(z) = \frac{4}{\sqrt{\pi}} \frac{z^2}{(2l_{mn}^2)^{3/2}} \mathrm{e}^{-z^2/(2l_{mn}^2)} \tag{S23}$$

with

$$l_{mn}^2 = \frac{b^2}{3} \frac{mn}{m+n}. \tag{S24}$$

This case happens with the probability $p_\infty^2(1-p_\infty)^{m+n-2}$. Averaging eq. (S23) with respect to $m$ and $n$ leads to the form

$$\psi(z) = \sum_{n=1}^{\infty} \sum_{m=1}^{\infty} p_\infty^2 (1-p_\infty)^{m+n-2} \psi_{m,n}(z). \tag{S25}$$

To derive eq. (S25), we assumed that $(1-p_\infty)^{m+n-2}$ decreases fast enough so that one can replace the upper bound of the sums with respect to $m$ and $n$ to the infinity.

The kinetics equation of the binding and unbinding of the $l$-th unit has the form

$$\frac{dp_\infty}{dt} = k_{\rm on}\Psi(p_\infty)(1-p_\infty) - k_{\rm off}p_\infty, \tag{S26}$$

where $\Psi$ is the probability with which the $l$-th unit is located in $0 < z < b$ and has the form

$$\Psi(p_\infty) = \int_0^b dz\, \psi(z). \tag{S27}$$

By solving eq. (S26) for the steady state, $dp_\infty/dt = 0$, the binding probability $p_\infty$ is derived as

$$\frac{k_{\rm off}}{k_{\rm on}} = \frac{(1-p_\infty)}{p_\infty}\Psi(p_\infty), \tag{S28}$$

see the black broken line in fig. S3. The right side is a function only of the binding probability $p_\infty$ and the left side $k_{\rm off}/k_{\rm on}$ is the binding constant. Eq. (S28) agrees with the prediction of the Rouse dynamics simulation for cases in which the number $N$ of adhesive units is large and the binding constant $k_{\rm off}/k_{\rm on}$ is small, see the dots and the black broken line in fig. S3.

#### S1.4 Extension to adhesive section of finite length

Eq. (S28) is simpler than the Rouse dynamics simulations, but its applicability is limited, see fig. S3. Eq. (S28) is derived by assuming that the number of units in the adhesive section is infinite, while the number of units in the adhesive section is finite in the Rouse dynamics simulation. The upper bound of the number of units bound to the surface and the effect of the ends of the adhesive section are neglected in the self-consistent scheme. We here take into account the upper bound of the number of bound units in an extension of the infinite approximation.

To take into account the upper bound of the number of bound adhesive units, we analyze the kinetics of the state transition, where the state is determined by the number $n$ of adhesive units, see fig. S4. For cases in which at least one unit is bound to the surface, $n > 1$, the number of bound units increases with the rate $\tilde{k}_{\text{on}}$ and decreases with the rate $\tilde{k}_{\text{off}}$. In general, the rates $\tilde{k}_{\text{on}}$ and $\tilde{k}_{\text{off}}$ are different from the rates $k_{\text{on}}$ and $k_{\text{off}}$ because of the connectivity of the units. The rates $\tilde{k}_{\text{on}}$ and $\tilde{k}_{\text{off}}$ do not depend on the choice of units because we still neglect the end effect. The kinetics of the probability $P_n$ that $n$ units are bound to the surface has the form

$$\begin{aligned} \frac{d}{dt}P_n(t) &= (N-(n-1))\tilde{k}_{\text{on}}P_{n-1} + \tilde{k}_{\text{off}}(n+1)P_{n+1} \\ &\quad -(N-n)\tilde{k}_{\text{on}}P_n - \tilde{k}_{\text{off}}nP_n \end{aligned} \tag{S29}$$

for $2 < n < N-1$ and

$$\frac{d}{dt}P_N(t) = \tilde{k}_{\text{on}}P_{N-1} - \tilde{k}_{\text{off}}NP_N \tag{S30}$$

for $n = N$ because of the upper bound of the number of bound units. In the steady state, $dP_n/dt = 0$, the probability $P_n$ has the form

$$P_n = \frac{(N-1)!}{n!(N-n)!}\left(\frac{p_\infty}{1-p_\infty}\right)^{n-1} P_1 \tag{S31}$$

for $n \leq 1$. We used

$$p_\infty = \frac{\tilde{k}_{\text{on}}}{\tilde{k}_{\text{on}} + \tilde{k}_{\text{off}}} \tag{S32}$$

in the expression of eq. (S31) (many readers would agree that eq. (S32) is apparent, but if one wants to prove it only by mathematical operations, one should do the following calculation without using eq. (S32) and find $p_\infty$ by taking the limit $N \to \infty$ to eq. (S35)).

The probability $q_{\rm on}$ that at least one unit is bound to the surface has the form

$$
\begin{aligned}
q_{\rm on} &= \sum_{n=1}^{N} P_n \\
&= \frac{P_1}{N}\frac{1-p_\infty}{p_\infty}\left(\frac{1}{(1-p_\infty)^N}-1\right).
\end{aligned}
\tag{S33}
$$

The ratio $p_{\rm on}$ of bound units has the form

$$
\begin{aligned}
p_{\rm on} &= \frac{1}{N}\sum_{n=1}^{N} nP_n \\
&= \frac{P_1}{N}\frac{1}{(1-p_\infty)^{N-1}}
\end{aligned}
\tag{S34}
$$

The (conditional) probability $p$ that an arbitrary unit is bound to the surface in the condition that at least one unit is bound to the surface thus has the form

$$
p = \frac{p_{\rm on}}{q_{\rm on}} = \frac{p_\infty}{1-(1-p_\infty)^N}. \tag{S35}
$$

Eq. (S35) returns to $p = p_\infty$ for $N \to \infty$ and to $p = 1$ for $N = 1$. Eq. (S35) greatly improves the infinite approximation, see the dots and solid lines in fig. S3.

#### S1.5 Scaling theory

The right side of eq. (S28) is a complex function of $p_\infty$. We here derive a simple approximate form of $p_\infty$ by using the scaling argument by de Gennes.[3] The number of units bound to the surface is $p_\infty N$ and thus, on average, every $p_\infty^{-1}$ units are bound to the surface. The size of a subsection between neighboring bound units is $\xi_{\rm i} = bp_\infty^{-1/2}$ if the subsection is a ideal chain. Each unit is thus confined in a layer of thickness $\xi_{\rm i}$ at the surface, see fig. S1**c**. The kinetic equation of the binding and unbinding of units thus has the form

$$
\frac{d}{dt}p_\infty = k_{\rm on}\frac{b}{\xi_{\rm i}}(1-p_\infty) - k_{\rm off}p_\infty, \tag{S36}
$$

where $b/\xi$ in the first term represents the fact that a unit is confined in a layer of thickness $\xi_{\rm i}$. In the steady state, $dp_\infty/dt = 0$, eq. (S36) has the solution

$$\frac{k_{\rm off}}{k_{\rm on}} = \frac{1 - p_\infty}{\sqrt{p_\infty}}. \tag{S37}$$

Eq. (S37) can be even solved in terms of $p_\infty$ as

$$p_\infty = 1 + \frac{1}{2}\left(\frac{k_{\rm off}}{k_{\rm on}}\right)^2 - \sqrt{\frac{1}{4}\left(\frac{k_{\rm off}}{k_{\rm on}}\right)^4 + \left(\frac{k_{\rm off}}{k_{\rm on}}\right)^2}. \tag{S38}$$

Numerically, eq. (S37) agrees well with eq. (S28), see fig. S5. We thus use this scaling theory to take into account the polymeric nature of tandemly repeated genes in the binding of nascent RNAs to RDRC/Dicers at the surface of the nulcear envelope and use eq. (S35) to take into account the fact that the number of genes in the repeat is finite. Fig. 1**c** in the main article is derived by substituting eqs. (S38) into eq. (S35) ($p$ is the fraction of bound units, provided that at least one unit is bound to the surface).

#### S1.6 Binding probability

The probability $q_{\rm on}$ $(= 1 - P_0)$ that more than one unit is bound to the surface is derived by the kinetic equation

$$\frac{d}{dt}P_0 = -k_{\rm on}N\Psi P_0 + k_{\rm off}P_1, \tag{S39}$$

see fig. S4. For the case of $N \ll N_{\rm L}$, $\Psi$ is approximated to eq. (S2. In the steady state, $P_1$ has the form

$$P_1 = \frac{k_{\rm on}}{k_{\rm off}}N\Psi(1 - q_{\rm on}). \tag{S40}$$

By using eq. (S33), the binding probability $q_{\rm on}$ has the form

$$q_{\rm on} = \frac{1}{1 + \frac{k_{\rm off}}{k_{\rm on}} \Psi^{-1} \frac{p_\infty (1-p_\infty)^{N-1}}{1-(1-p_\infty)^N}}. \tag{S41}$$

Fig. 1**b** in the main article is derived by substituting eq. (S38) into eq. (S41).

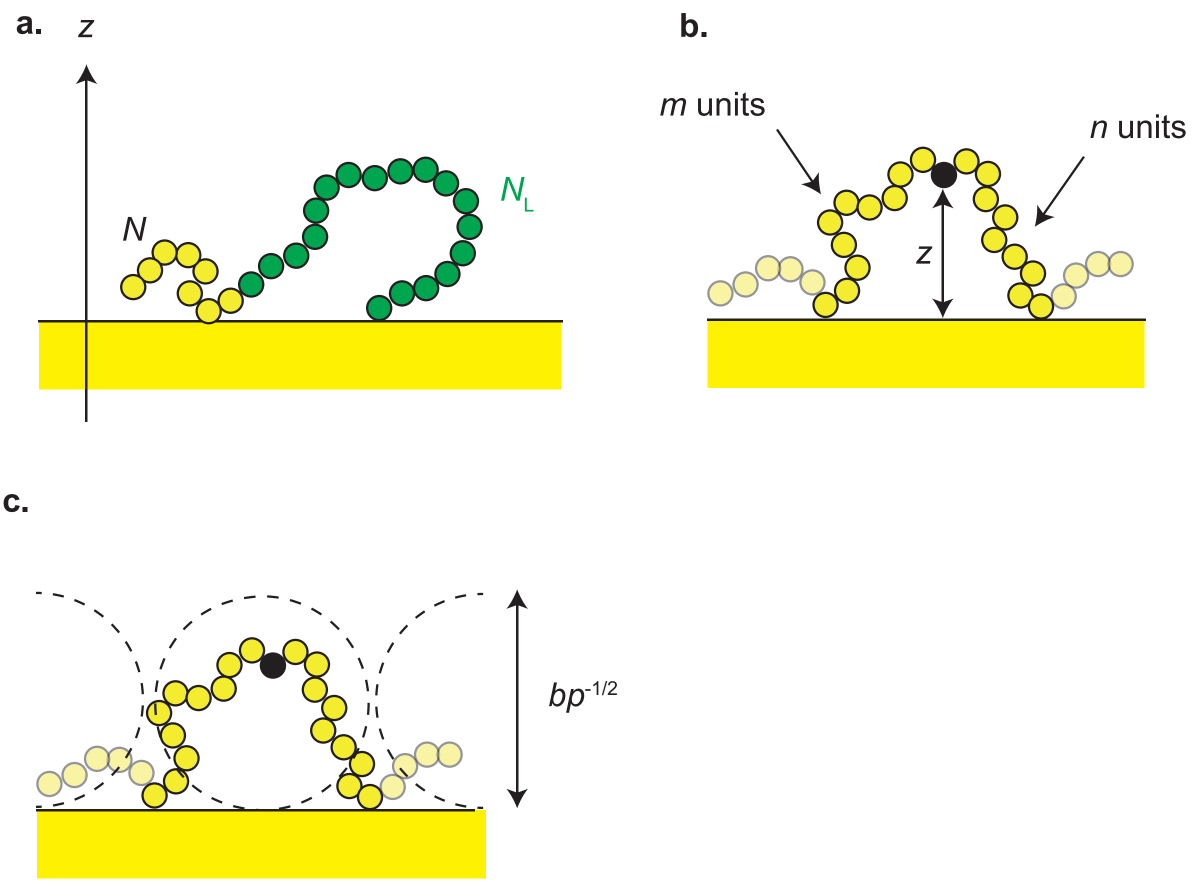


Figure S1: **Model of polymer surface adhesion :** Models to analyze the adhesion of polymers to a surface by simulation (**a**), self-consistent scheme (**b**), and scaling theory (**c**). The adhesive units are shown by yellow beads and the linker units are shown by green beads. The positions of the beads is represented by the distance from the surface.

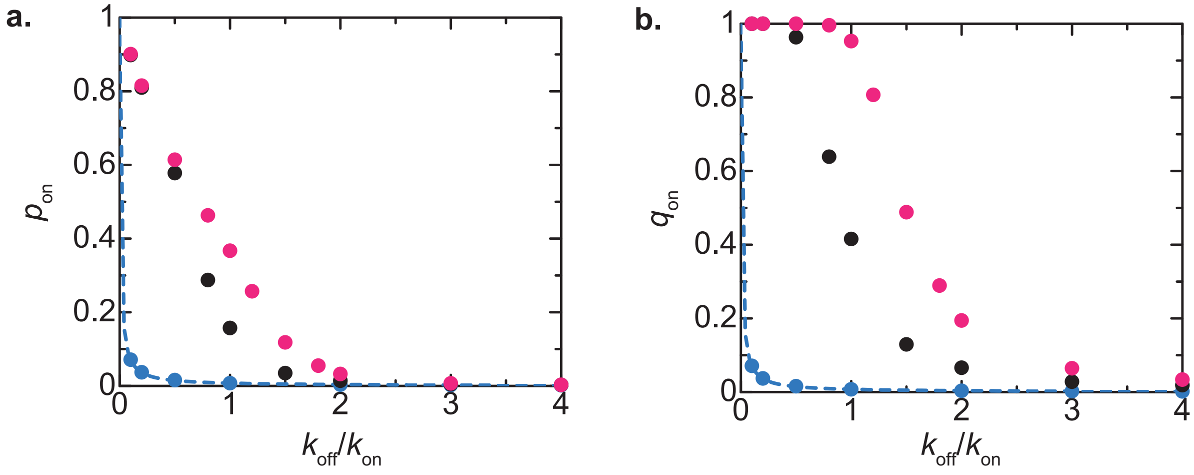


Figure S2: **Binding probability derived by simulations:** The ratio $p_{\rm on}$ of bound units and the probability $q_{\rm on}$ that more than one adhesive unit in a polymer is bound to the surface are shown as functions of the binding constant $k_{\rm off}/k_{\rm on}$ for $N = 1$ (cyan), 10 (black), and 20 (magenta). The dots are derived by the Rouse dynamics simulation and the cyan broken curve is derived by using eq. (S4). The values of parameters used in the simulations are summarized in Table S1.

Table S1: Values of parameters used in the simulation.

| Symbol | Meaning | Value |
|---|---|---|
| $\tau\xi$ | Monomer relaxation time | 10.0 |
| $N_{\rm L}$ | Linker length | 200 |
| $z_{\rm m}/b$ | Width of surface-monomer interaction potential | 0.3 |
| $\xi\Delta t$ | Time step | 0.1 |
| $M$ | Number of trials | 40 |
| $\xi t$ | Simulation time | $> 4.0 \times 10^6$ |
| $\xi t_{\rm eq}$ | Equilibriation time | $> 4.0 \times 10^6$ |

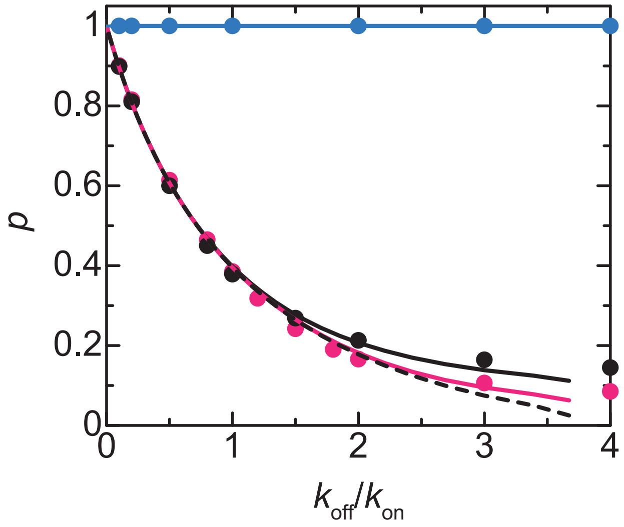


Figure S3: **Binding probability derived by self-consistent scheme:** The conditional probability $p$ that an arbitrary adhesive unit in a polymer to the surface, provided that at least one adhesive unit is bound to the surface, is shown as a function of the binding constant $k_{\rm off}/k_{\rm on}$ for $N = 1$ (cyan), 10 (black), 20 (magenta). The dots are derived by using $p = p_{\rm on}/q_{\rm on}$ to the data obtained by the Rouse dynamics simulation (see also fig. S2). The black broken line is derived by using eq. (S28). The solid lines are derived by using eq. (S28) to eq. (S35).

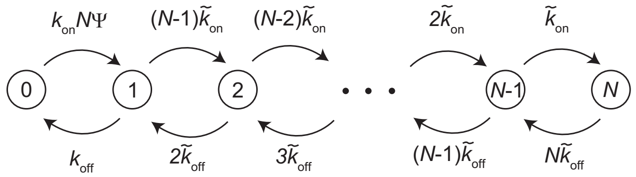


Figure S4: **Diagram of state transitions:** We analyze the state transition to take into account the upper bound of the number of adhesive units bound to the surface. The states are represented by the number $n$ of adhesive units bound to the surface (shown by the number in the circle). For $n > 1$, the number of adhesive units increases with the rate $\tilde{k}_{on}$ and decreases with the rate $\tilde{k}_{off}$. $\tilde{k}_{on}$ and $\tilde{k}_{off}$ are, in general, different from $k_{on}$ and $k_{off}$ because of the connectivity of the adhesive units and do not depend on adhesive units because we still neglect the end effect.

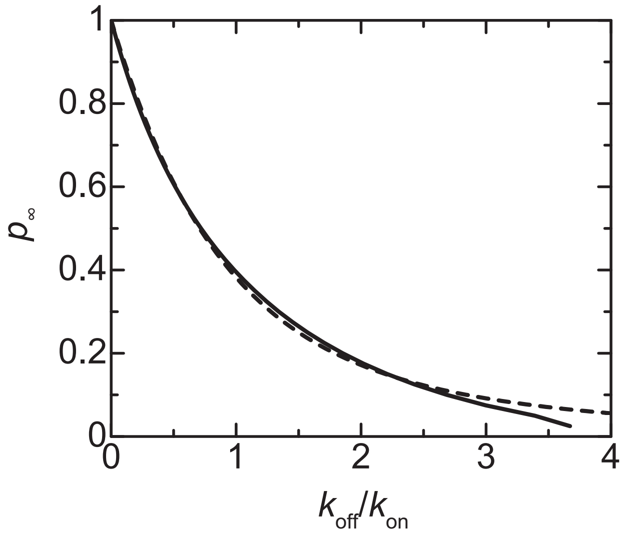


Figure S5: **Binding probability derived by using the scaling theory:** The probability $p_\infty$ that an arbitrary unit in a long chain is bound to the surface is shown as a function of the binding constant $k_{\text{off}}/k_{\text{on}}$. The solid line is derived by using the self-consistent scheme, see eq. (S28), and the broken line is derived by using the scaling theory, see eq. (S37).
